# A patient derived missense mouse model of Kabuki syndrome 1

**DOI:** 10.1101/2025.11.13.688040

**Authors:** Sara Tholl Halldorsdottir, Meghna Vinod, Hilmar Orn Gunnlaugsson, Ellen Dagmar Bjornsdottir, Teresa Luperchio, Jill A. Fahrner, Agnes Ulfig, Hans Tomas Bjornsson

## Abstract

Kabuki syndrome (KS) is a rare cause of intellectual disability resulting from heterozygous pathogenic variants in the gene encoding the histone methyltransferase KMT2D. A previously established loss-of-function mouse model of KS exhibits key phenotypic features, and therapeutic trials in this mouse model suggest postnatal malleability of neurological symptoms. However, 15-30% of individuals with KS, carry missense variants. To investigate whether missense variants lead to similar phenotypic presentation in mice, we used CRISPR-Cas9 to introduce the KS patient variant R5230H into C57BL/6NTac. Computational and *in vitro* testing suggests that the R5230H variant does not impair protein stability or loss of enzyme function of KMT2D. Despite a distinct mechanistic basis, our new mouse model (*Kmt2d^+/R5230H^*) recapitulates most phenotypes of our prior loss-of-function model, including growth deficiency, craniofacial anomalies, and IgA deficiency but not altered neurological function. *Kmt2d^+/R5230H^*mice show perinatal lethality and a high frequency of unilateral kidney agenesis, a novel phenotype in KS mouse models. *Kmt2d^+/R5230H^* mice provide a unique opportunity to understand the impact of missense variants on KMT2D function and uncover developmental and perinatal abnormalities in KS.

**Summary statement:** A novel Kabuki syndrome missense mouse model with intact KMT2D enzymatic function shares most features with prior KS models, except disruption of adult neurogenesis, and exhibits novel unilateral kidney loss.

## Introduction

Extensive work has been conducted over the last 10 years on therapeutic development for Kabuki syndrome (KS) (Benjamin et al., 2017; Bjornsson et al., 2014; L. Zhang et al., 2021). This work has been conducted using a mouse model (*Kmt2d^+/βGeo^*) which shows many defining features of individuals with KS (Comel et al., 2025; Harris et al., 2019; Ruault et al., 2020; Schott et al., 2017). These include growth deficiency with flattened facial profile (Bjornsson et al., 2014; Fahrner et al., 2019), a visuospatial defect (Bjornsson et al., 2014) and IgA deficiency (Pilarowski et al., 2020). Although, this model is thought to have a similar loss of function mutational mechanism to the majority of individuals with KS (Hannibal et al., 2011), it was, however, generated by inserting an artificial β-galactosidase neomycin cassette which theoretically could have additional effect on the function of the KMT2D protein such as altered structure resulting in aberrant binding. Moreover, this technically only reflect a single loss of function allele and the molecular cause for about 15-30% of KS individuals involves heterozygous missense variants (Cocciadiferro et al., 2018; Hannibal et al., 2011), highlighting that this model is not representative of the entire KS population. These KS causing missense variants have been shown to be enriched at the enzymatic (SET) and the FYRN domains (Faundes et al., 2019). In contrast, Wiedemann-Steiner syndrome, caused by variants in a gene encoding another related H3K4 methyltransferase, KMT2A, shows enrichment at the CpG binding domain (CXXC) but not the enzymatic SET domain (Reynisdottir et al., 2022), suggesting that the mechanism of disease may be distinct despite overlapping H3K4-methyltransferase function. These observations highlight the importance of the binding domains (CXXC, FYRN) in these two methyltransferases, KMT2A and KMT2D.

Multiple treatment strategies countering the observed epigenetic abnormalities have shown promise in a mouse model rescuing the neurological deficits in postnatal life (Benjamin et al., 2017; Bjornsson et al., 2014; L. Zhang et al., 2021). However, it remains unclear whether the same treatments would also work in mouse models with a different mutational mechanism. One prior model harboring a missense variant exists which is associated with a KS-like phenotype (Yamamoto et al., 2019). This missense variant lies within exon 13 and functions as a hypomorph with phenotypes only observed in homozygous animals which does not reflect what is seen in individuals with KS.

Here, we use CRISPR-Cas9 to introduce a patient-derived variant into the corresponding sequence of the murine *Kmt2d*. Although protein stability and enzyme function of KMT2D is not affected, *Kmt2d^+/R5230H^* mice have extensive perinatal lethality, and many overlapping phenotypes with the prior model including a craniofacial defect, growth deficiency, IgA deficiency and fewer Peyer’s patches. However, phenotypes of the altered neurological function and visuospatial defect or decreased global histone methylation levels seen in the prior model were not found in these mice. In addition, kidney agenesis emerged as a novel phenotype that has not been previously observed in KS mouse models. We believe that this model will be useful for the KS community to aid in the investigation of the mechanistic basis of KS and to explore both previous and novel therapeutics modalities for KS.

## Results

### High mortality in a mouse model with a novel missense variant in the FYRN domain

In an effort to extend the availability of representative KS1 mouse models we aimed to generate a mouse model with a patient specific missense variant. We examined the list of patients with missense variants seen in our Epigenetics and Chromatin Clinic at Johns Hopkins Hospital. We selected a variant (c.G15536A:p.R5179H), located within the FYRN domain and seen in one of our patients with classical symptoms of KS and a score comparable to what is seen in patients with loss-of-function variants (**Table S1**). Mice carrying the corresponding mutation (*Kmt2d^+/R5230H^*) were generated using CRISPR-Cas9 and subsequently subjected to phenotyping. The corresponding mouse variant involves a change from an arginine (R) to a histidine (H) at a conserved region within the FYRN domain (c.15689_90GA>AC:p.R5230H; **Figure 1A-B**). CRISPR-Cas9 off-target effects were excluded at the predicted *Samd8* locus by screening F1 animals and F4 offsprings (**Figure S1A**). Additionally, the mouse phenotypes were examined throughout five generations since a second variant would be expected to randomly segregate off and therefor unlikely to be the cause of the observed phenotypes. We used AlphaFold3 to model the potential structure of the FYRN/C domain and the change in structure upon introduction of R5230H. AlphaFold predicts (>70% confidence) that this variant localizes within a β-sheet and that the change will not affect the structure of KMT2D (**Figure 1C**). Visualization of the predicted electrostatic potential of this domain however displays a change to a less positive charge (**Figure 1D**). To investigate potential interactors of this region we performed protein-protein prediction against components of the COMPASS complex using a part of the C-terminal region of KMT2D. Aside from WDR5 that is known to bind to the WIN motif at this region, no interactions were predicted with high confidence (ipTM>0.8). However, when visualized, two predictions resulted in a probable interaction shown with connectors to the specific region of the variant, NCOA6 (**Figure S1B**) and/or the PAXI1-PGR1A partners (**Figure S1D**). In both instances, this interaction was predicted to be present at the amino acid R5230 and not predicted after the R5230H substitution (**Figure S1C and S1D**).

**Figure 1.**
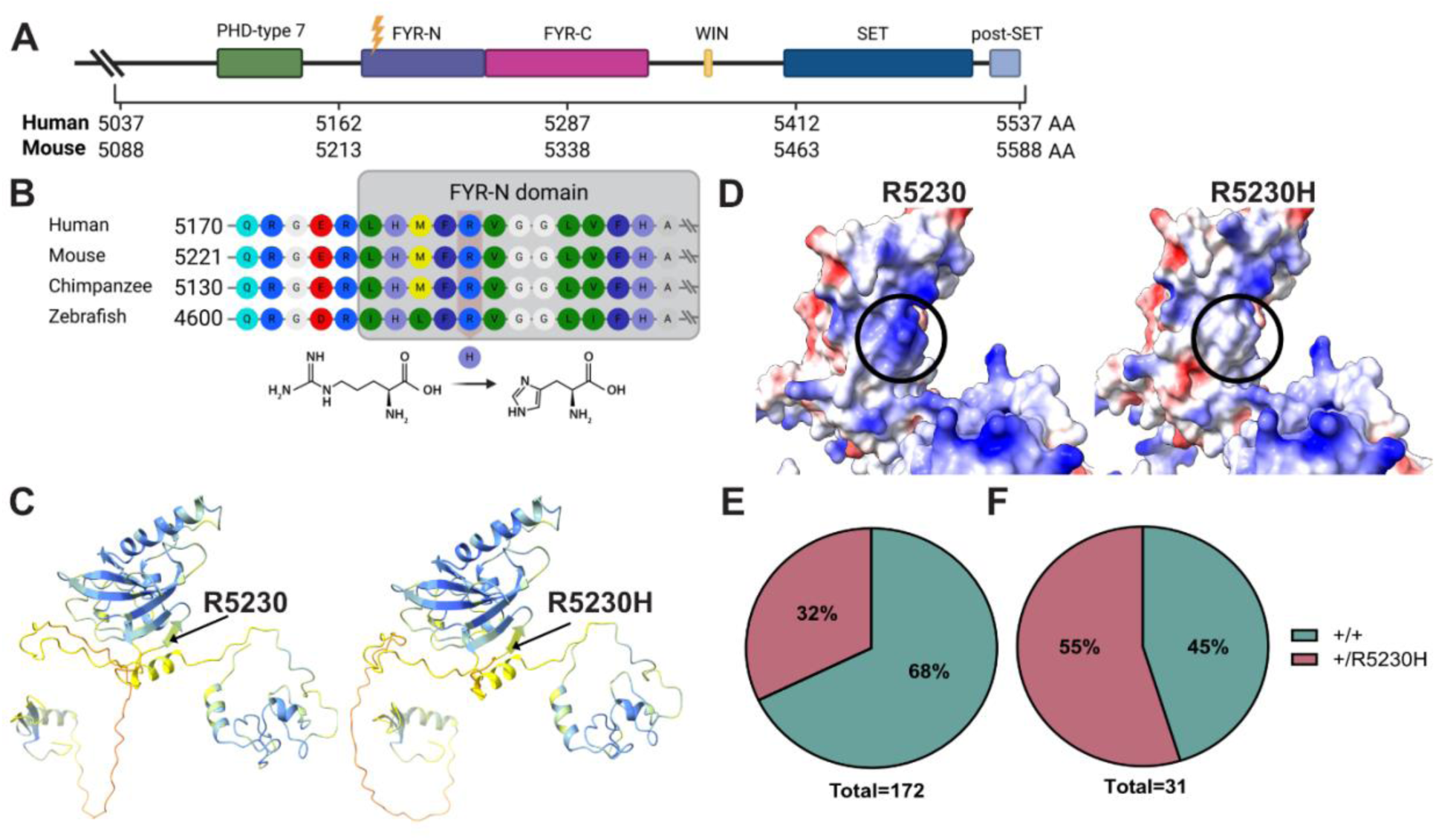
The R5230H variant changes electrostatic charge but not the structure of the FYRN domain. (A) A schematic view of the human and mouse KMT2D C-terminus including 500 amino acids with variant location marked with a lightning symbol. (B) Partial amino acid sequences around the missense variant in various organisms with the R5230H substitution highlighted in addition to the molecular structures of arginine and histidine. Amino acids colored by the RasMol amino acids color scheme. AlphaFold3 prediction for FYRN/C domain with 100 amino acids flanking (C) colored by pLDDT with orange representing the least confidence and dark blue representing the highest confidence or by (D) electrostatic charge with amino acid R5230 circled. Blue color indicates a positive charge and red color, a negative charge. Mendelian ratios for (E) postnatal and (F) embryonic (E16.5-18.5) survival. Total indicates the number of animals used for calculations.

Similar to what is seen in *Kmt2d^+/βGeo^* mice, we observed homozygous lethality of mice carrying the variant (data not shown). In contrast to the previous model, however, in the heterozygous state, the Mendelian ratio was much more skewed (Bjornsson et al., 2014), with only ∼30% live-born heterozygotes (50% expected) (**Figure 1E**). This number does, however, not consider cohorts with no live born heterozygotes, indicating an even higher perinatal lethality of *Kmt2d^+/R5230H^* compared to *Kmt2d^+/βGeo^*mice. To investigate if the lethality occurred during embryonic period, we performed timed breeding and sacrificed dams at E16.5-18.5. In those cases, we observe a Mendelian ratio closer to what is expected with 55% of the embryos carrying the R5230H variant (**Figure 1F**). This suggests that lethality may occur perinatally rather than during embryonic life.

### The R5230H variant does not alter expression or enzymatic function of KMT2D

Gene expression studies on hippocampal tissue showed that the R5230H variant does not affect stability of *Kmt2d* mRNA, as evident by unaltered *Kmt2d* mRNA levels between *Kmt2d^+/R5230H^* and *Kmt2d^+/+^*mice when amplifying either regions upstream (exon 3-4 and 4-5) or downstream (exon 49-50 and 51-52) of the variant (**Figure 2A**). Similarly, a Western blot using spleen tissue for KMT2D did not show any significant differences in the protein levels between samples derived from *Kmt2d^+/R5230H^* and *Kmt2d^+/+^* animals (**Figure 2B**). When testing the global enzymatic activity by Western blot, we did not observe decreased H3K4me1 or H3K4me3 levels in *Kmt2d^+/R5230H^* compared to *Kmt2d^+/+^*mice (**Figure 2C-D**). Together, these results suggest that the R5230H variant does not affect KMT2D protein stability or cause a drastic loss of KMT2D enzymatic function.

**Figure 2.**
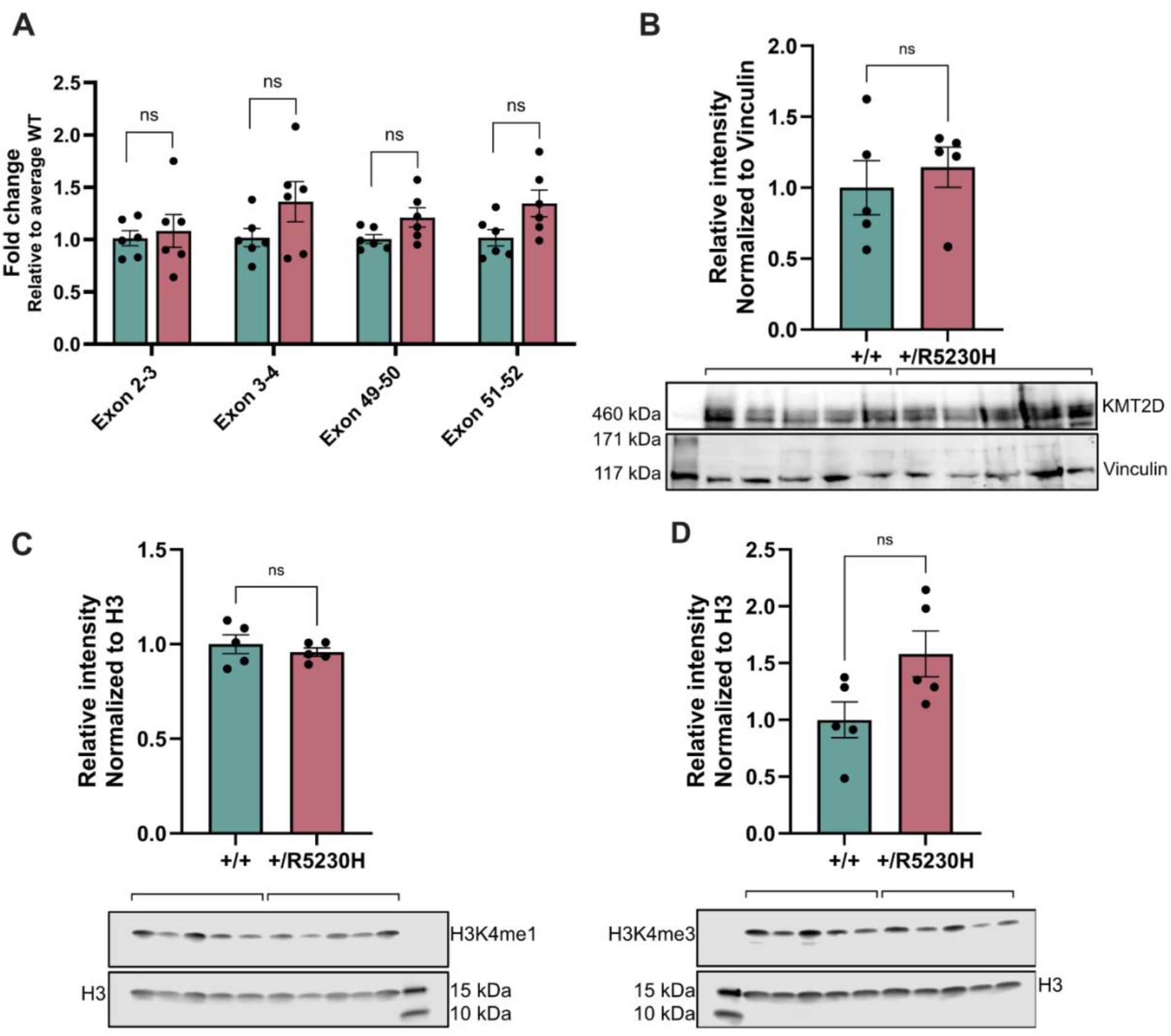
R5230H does not cause changes in KMT2D stability or global methylation. (A) Relative mRNA expression of *Kmt2d* from hippocampal tissue detected with four primer pairs (exon 2-3, 3-4, 49-50 or 51-52). Western blot and quantification of (B) KMT2D with Vinculin as control, (C) H3K4me1 or (D) H3K4me3 using H3 as control with protein isolated from spleen tissue. Both males and females were used (n=5-6 per genotype). Data is represented as mean ± SEM. Statistical tests were performed with an unpaired Student’s t tests.

### *Kmt2d^+/R5230H^* mice show core phenotypes of KS1 except disruption of adult neurogenesis

Similar to *Kmt2d^+/βGeo^* mice (Fahrner et al., 2019), *Kmt2d^+/R5230H^* mice show growth deficiency with significantly decreased body weight compared to *Kmt2d^+/+^* mice (P<0.0001; **Figure 3A**). We performed CT scans of the skulls to assess the craniofacial structure and measured the angle of the neurocranium by using 3 landmarks: posterior of the skull to bregma and the border of frontonasal suture (**Figure 3B**). With these measurements we observed a sharper angle in *Kmt2d^+/R5230H^* mice (**Figure 3C**) indicating more arching of the cranium similar to *Kmt2d^+/βGeo^* mice (Fahrner et al., 2019). Other features that resembled the previous model were the significantly decreased body length (**Figure S2A**), decreased long bone length (Femur; P<0.01, Tibia; P<0.05; **Figure 3D**), lower IgA blood serum levels (P<0.0001; **Figure 3E**) and reduced number of Peyer’s patches (P<0.01; **Figure 3F**) compared to *Kmt2d^+/+^*mice. These results suggest that these phenotypes are not dependent on the type of variant or secondary to the βGeo tag in the prior model. Behavioral testing showed that, similar to *Kmt2d^+/βGeo^* mice, *Kmt2d^+/R5230H^* mice show no significant difference in open field testing (**Figure S2B**) compared to *Kmt2d^+/+^* mice or decreased distance traveled in a Morris water maze (MWM, **Figure 4A**) indicating that they do not display anxiety-like behavior. However, unlike the *Kmt2d^+/βGeo^* mice, *Kmt2d^+/R5230H^* mice show no significant difference in Morris water maze platform crossings (**Figure 4B)** or proportion of time spent in the platform quadrant (**Figure 4C**) compared to littermates (i.e., *Kmt2d^+/+^* mice). To further test the visuospatial memory of these mice, we also performed a Y-maze. Similar to the MWM, this behavior test showed no significant difference in novel arm entry (**Figure 4D**), time spent in the novel arm (**Figure 4E**) or spontaneous alternations (**Figure 4F**) compared to *Kmt2d^+/+^*animals. To further validate unaltered neurodevelopment on a cellular level, we stained cryosectioned brains and examined the region of the dentate gyrus **(Figure 4G)**. Contrary to what was seen in *Kmt2d^+/βGeo^* mice, we did not observe a reduced volume of the dentate gyrus granule cell layer (**Figure 4H**) or a difference in numbers of 5-ethynyl-2′deoxyuridine (EdU) stained cells in the hippocampus **(Figure 4I)** suggesting relatively normal proliferation or differentiation of neurons of the hippocampus. Additionally, staining of the hippocampus for H3K4me3 did not reveal decreased methylation levels (**Figure 4J**) as seen in our previous *Kmt2d^+/βGeo^*mouse model. This supports what we previously described for spleen tissue **(Figure 2D)**, i.e., unchanged global methylation levels upon the introduction of the R5230H variant. Together, these findings suggest that our novel mouses model replicates the majority of core disease phenotypes except features related to a disruption of adult neurogenesis, a phenotype that has previously been rescued with agents that affect the epigenetic machinery.

**Figure 3.**
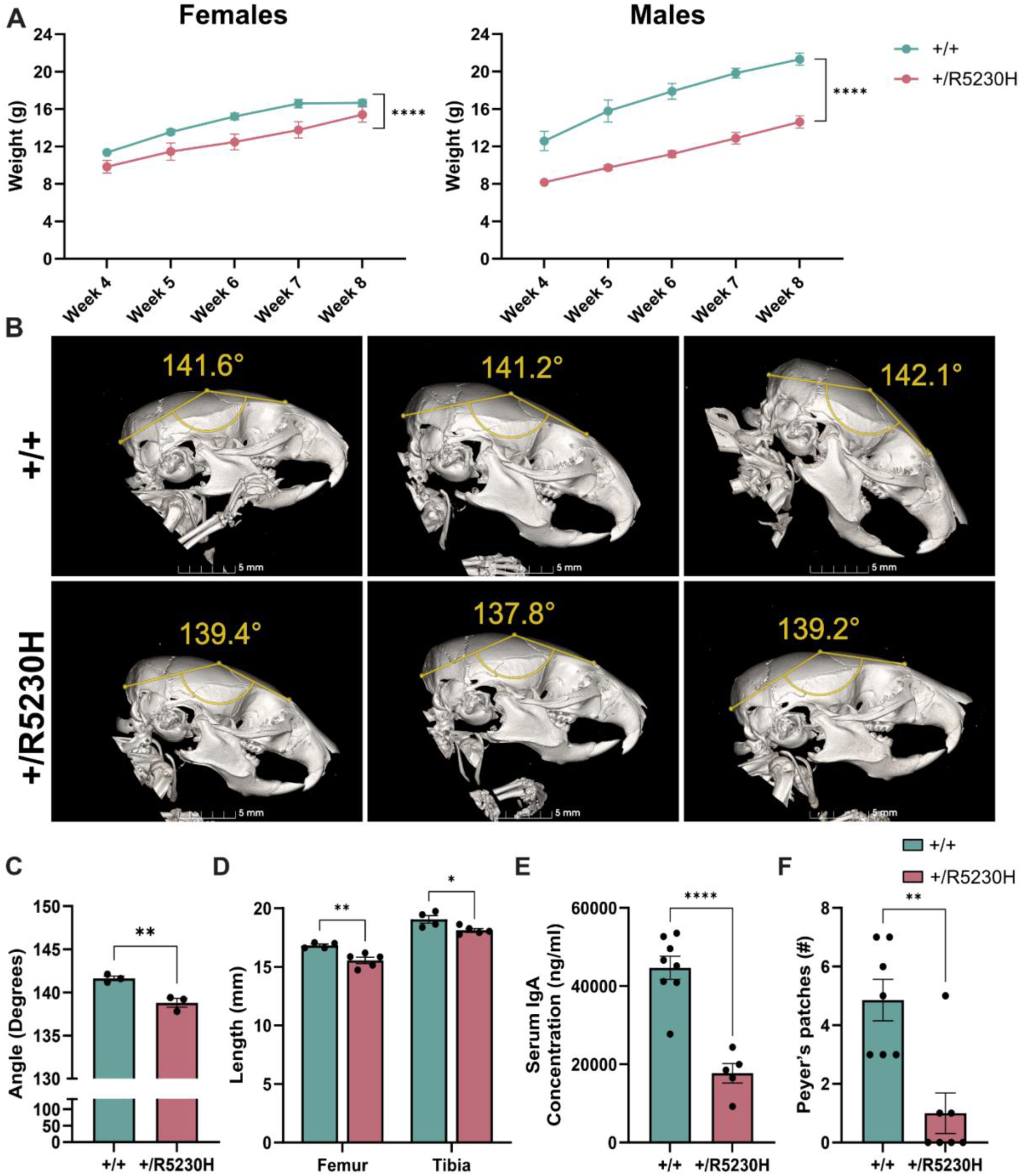
Concordance of phenotypes in *Kmt2d^+/R5230H^* and *Kmt2d^+/βGeo^* mouse models. (A) Body weights of the mice compared in both female (left, n=6-8) and male (right, n=6-13) mice. (B) Reconstruction of lateral view CT for *Kmt2d^+/+^* and *Kmt2d^+/R5230H^* craniofacial profiles with measured landmarks in yellow dots and angle shown as degrees (n=3). (C) Angle measurements for *Kmt2d^+/+^* and *Kmt2d^+/R5230H^*neurocranium (n=3). (D) Femur and tibia measurements of 3–4-month-old mice in mm (n=4-5). (E) Levels of IgA in blood serum quantified at 3 months of age. One outlier was detected among the *Kmt2d^+/R5230H^* (n = 6) using Grubbs’ test (p < 0.05) and was removed prior to analysis. The final sample size for *Kmt2d^+/R5230H^* was n=5 and *Kmt2d^+/+^* n = 8. (F) Number of Peyer’s patches quantified by counting (n=7). Data are represented as mean ± SEM. Statistical tests were performed with mixed-effects ANOVA (A) or unpaired Student’s t tests (C-F). *P ≤ 0.05, **P ≤ 0.01, ***P ≤ 0.001, ****P ≤ 0.0001.

**Figure 4.**
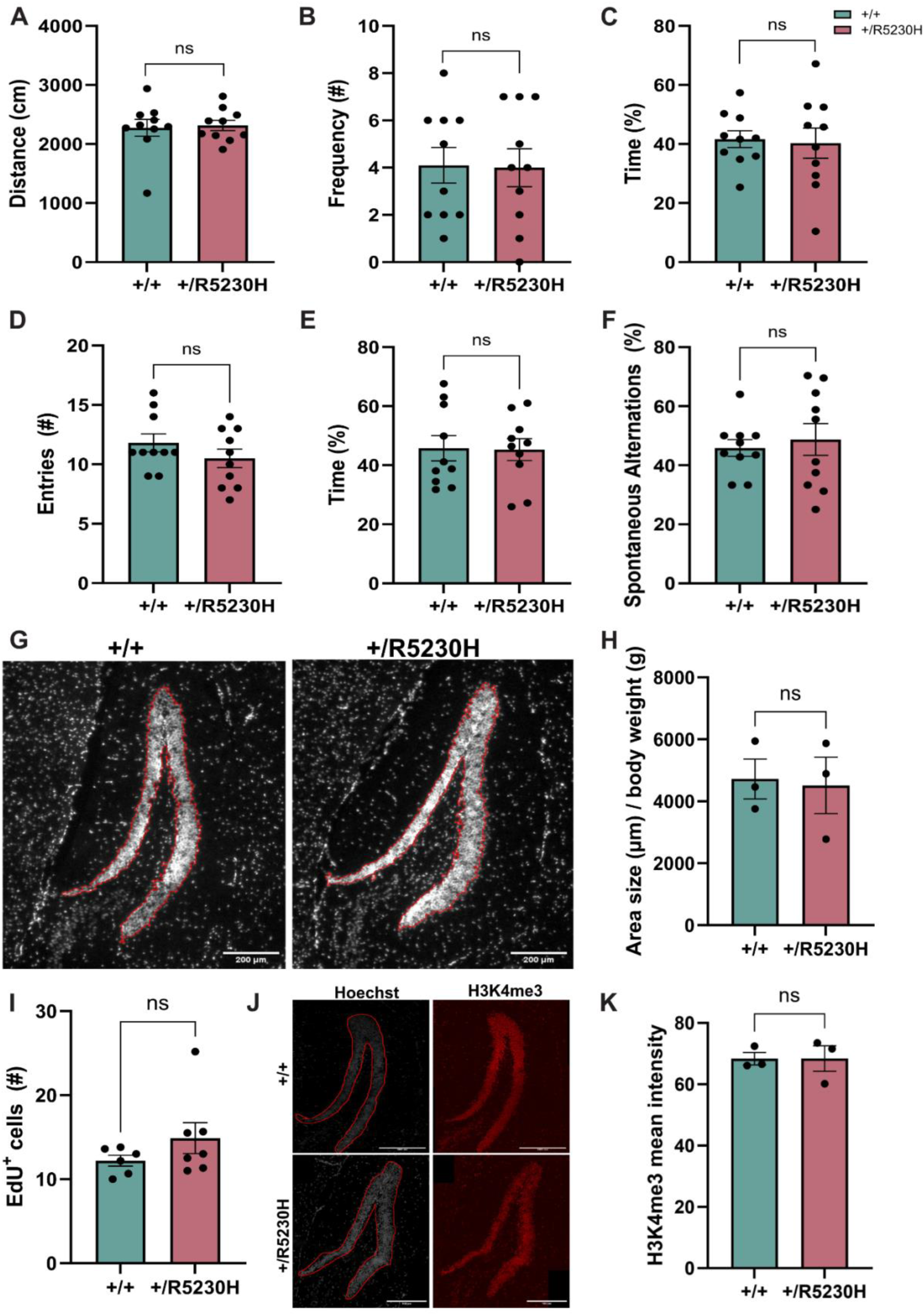
Discordance between *Kmt2d^+/R5230H^* and *Kmt2d^+/βGeo^*mouse models. Morris water maze analysis measuring (A) total distance traveled, (B) total platform crossings and (C) percentage of time spent in the platform quadrant relative to total time spent in maze. Y-maze analysis measuring (D) the number of novel arm entries, (E) the percentage of time spent in novel arm relative to total time and (F) spontaneous alternations. (G) Representative images of brain sections stained with the Hoechst dye with granule cell layer outlined and (H) quantification in µm. (I) Quantification of number of proliferative EdU+ cells within the granule cell layer. (J) Representative images of H3K4me3 staining within the granule cell layer. Hippocampus area measured was outlined with Hoechst staining. (K) Quantification of H3K4me1 hippocampal mean intensity per area. Data is represented as mean ± SEM. Statistical tests were performed with an unpaired Student’s t tests. This study was limited to females (n=10) since males were used for breeding.

### *Kmt2d^+/R5230H^* mice display novel phenotypes not seen in other KS mouse models

About 43% of *Kmt2d^+/R5230H^* animals show unilateral renal agenesis (**Figure 5A**). This abnormality was not seen in any of the *Kmt2d^+/+^* animals and has not been observed or reported in *Kmt2d^+/βGeo^* mice. Loss of a single kidney often results in enlargement of the remaining kidney (Woolf & Hillman, 2007) governing further histological investigation. Histology staining revealed normal gross histology (**Figure 5C**), but size quantification showed enlargement of the glomeruli within the single kidney compared to kidney histology from *Kmt2d^+/R5230H^* animals that presented with two kidneys or *Kmt2d^+/+^* animals (**Figure 5B**). These results suggest that this particular model may be valuable for studying the impact of KMT2D dysfunction in kidney development and abnormalities.

**Figure 5.**
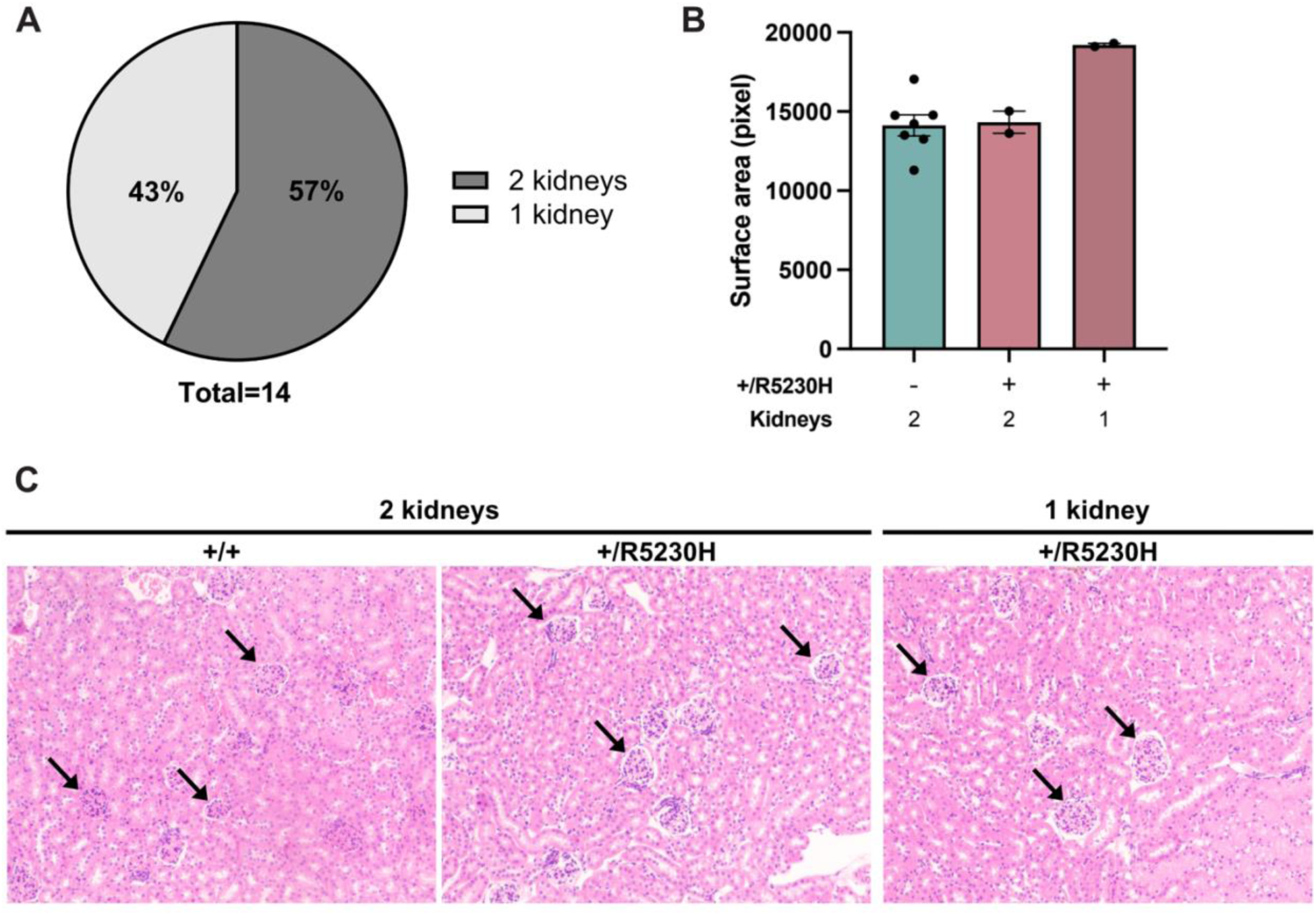
Novel phenotypes found in *Kmt2d^+/R5230H^* mice. (A) Number of kidneys observed in R5230H mice with number of animals shown as percentage. (B) Glomeruli size quantification and (C) representative images of H&E staining of kidneys from *Kmt2d^+/+^* (n=7) and *Kmt2d^+/R5230H^* mice with one (n=2) or two (n=2) kidneys. Data are represented as mean ± SEM.

## Discussion

Here, we describe a novel missense mouse model of KS based on a patient specific missense variant located in the FYRN domain of *KMT2D*. The FYRN domain is a known hotspot of missense variants specific to individuals with KS compared with variants found in the general population or cancers (Faundes et al., 2019). The selected patient variant is an arginine residue recurrently substituted in KS but similar recurrent substitutions have also been reported at Arg5048 and Arg5432 (Faundes et al., 2019). The FY-rich motifs N- and C-terminal form a single folded FYR domain enriched in phenylalanine/tyrosine that is found in many chromatin modifying proteins, specifically common in H3K4 methyltransferases (García-Alai et al., 2010). Limited information exists about the specific function of the FYR domain of KMT2D, but it has been shown that deletion of either part of the domain (FYRN or FYRC) in the demethylase, JMJ14, impairs its chromatin binding capacity rather than affecting its catalytic function (S. Zhang et al., 2015). Even though there are limitations in using AlphaFold for large proteins such as KMT2D, our predictions using about 400-500 amino acids of the C-terminal are consistent with previous reports, showing that the FYR domain adopts an α + β structure. Our predictions revealed that R5230H is located within a β-sheet which often mediates protein binding (García-Alai et al., 2010; Nowick, 2008) and suggests changes in electrostatic charge at predicted binding sites. The amino acid substitution is predicted to affect protein interaction of known components of the COMPASS, NCOA6 and the partners PAXI1-PGR1A, proteins that have both been implicated in the recruitment specificity of the complex (S. Lee et al., 2006).

The *Kmt2d^+/R5230H^* mice show concordance with the majority of phenotypes previously described in *Kmt2d^+/βGeo^* mice including growth deficiency, craniofacial defects, low IgA serum levels and decreased number of Peyer’s patches. Interestingly, the postnatal severity of the *Kmt2d^+/R5230H^* model appears to be greater than what is seen for *Kmt2d^+/βGeo^* mice as evaluated by the number of surviving heterozygotes. We also observe unilateral kidney agenesis but previous studies report that individuals with KS have up to 40% rate of renal malformations. Such abnormalities have not been described in earlier mouse models of KS and our findings therefore highlight a clinically relevant phenotype that can now be modeled and investigated in mice. The malformations found in individuals with KS have a wide range of presentations and severity which does not appear to correlate with variant type (Banka et al., 2012; Courcet et al., 2013; Hannibal et al., 2011). Similar to what we observe in our model, a prior case report showed unilateral renal agenesis with renal insufficiency in a patient with a truncating c.9829C>T, p.Gln3277X variant in *KMT2D* (Courcet et al., 2013). Future work will have to see whether this novel phenotype remains stable or is lost with additional breeding.

Some discrepancies in the observed phenotypes between the two models are noted, as we did not detect changes in hippocampal size or visuospatial deficits seen in the prior model. Notably, these phenotypes may be secondary to the disruption of adult neurogenesis, leading us to speculate that our model does not exhibit the same extent of neurological dysfunction as the prior model. In support of this interpretation, the patient carrying this variant presented with a higher than expected cognitive abilities, IQ of 77 and only a mild form of intellectual disability without any report of visuospatial defects. One possible explanation of why we do not detect defects in this mouse model is that perhaps intellectual disability must reach at least a moderate or severe level to be reliably modeled in mice. Alternatively, the use of only female subjects in our behavioral assays may have influenced the results if there is a sex difference. Additionally, the quantification of H3K4me3 within the hippocampus revealed no significant difference in global methylation. This finding, combined with preserved adult neurogenesis and hippocampal structure, suggests that the loss of KMT2D catalytic function may be necessary for the manifestation of the adult neurogenesis disruption in KS.

Genotype-phenotype studies have not found a definitive correlation between disease appearance and type of variants (Hannibal et al., 2011; Miyake et al., 2013). A study from 2012, exploring KMT2D missense variants suggests that perhaps not all KS-causing variants are loss-of-function (Banka et al., 2012). Concordantly, another more recent study tested missense variants from KS individuals *in vitro* and found that only about 65% appeared to reduce H3K4 methyltransferase activity (Cocciadiferro et al., 2018). Furthermore, a new disease entity has recently been described, where missense variants in exon 38 and 39 of *KMT2D* cause no alterations in histone methylation and the observed phenotypes are somewhat distinct from KS. In addition, these individuals did not present with intellectual disability (Cuvertino et al., 2020). Since we do not see changes in H3K4 methylation, this highlights an alternative function for KMT2D and the idea that a missense variant could cause changes in a function of KMT2D that is unrelated to its catalytic activity. KMT2D has previously been shown to be important during embryonic development where targeted deletion in cardiac precursors causes embryonic lethality (Ang et al., 2016) and whole-body homozygous deletion results in lethality around E9.5 (J. E. Lee et al., 2013). Although numerous studies highlight the critical role of KMT2D in development, and some propose a potential mechanistic basis, none have definitively explained its precise function (Froimchuk et al., 2017; Shan et al., 2024). Our findings demonstrate an essential role of KMT2D in development that may be independent of its methyltransferase function.

In summary, here we present the first patient-specific mouse model for KS, capturing core KS phenotypes along with high perinatal lethality and renal agenesis, making it a valuable tool for studying these phenomena in the future. Despite harboring the majority of previously observed KS features, it lacks defects in adult neurogenesis and loss of enzyme activity. We therefore speculate that KMT2D’s catalytic role might underly visuospatial defects and neurogenesis disruptions but no other KS traits, however, additional studies will be needed to robustly test this idea.

## Materials and methods

### Animals

#### Mouse model

The mouse model was created by Taconic with a constitutive knock-in of a point mutation via CRISPR-Cas9 with microinjection on C57BL/6NTac zygotes followed by in vitro fertilization (IVF). The targeting strategy was based on NCBI transcript NM_001033276.3 using gRNA (5’-CGGCTACA_CATGTTCCGAGT-AGG-3’) and homologous recombination donor (HDR) oligonucleotide containing the knock-in variant. Two founder mice were successfully established, and line was maintained on pure C57BL/NTac background. CRISPR-Cas9 off-targets were screened with Sanger sequencing. Polymerase chain reaction (PCR) was performed with primers targeting *Samd8* (5’-GCAGGTGTCGTTCTTATGTTGGGG-3’ and 5’-AGTCAGAGTCATTCCAAAGCCGC-3’) using ExtraMix which was provided by deCODE genetics and sequencing performed by deCODE genetics using two primers: 5’-ACAGTGACTATGTGAGGCAGGG-3’ or 5’-TTTGTGAGCTTGAGGTCAGCCTGG-3’. Genotyping was performed on ear clips with primers: R5230H F-(5’-CTTCAAGGACAAGACCATGCTG-3’) and R5230H R-(5’-AGGCAGGGGCATCATTTG-3’) targeting exon and intron 49, flanking the missense variant. The R5230H variant introduces a restriction site, making it possible to cleave the R2530H PCR product with BsaAI restriction enzyme. The following thermal cycler conditions were used for PCR: Initial denaturing for 2 min at 96°C followed by 40 cycles of denaturing at 96°C for 30 sec, annealing at 57°C for 30 sec and extension at 57°C for 1 min. Final extension was performed at 72°C for 10 minutes.

#### Housing

All animals were bred in a sterile facility at ArcticLAS ehf, under 12-hour light/dark cycles at 21-24°C and 40% humidity in cages of 6 mice maximum. Behavioral testing was done in a University Facility (VRIII). Mice were provided with filtered water and kept on separate diets for breeding (Altromin NIH#31 M) and maintenance (Altromin 1324) *ad libitum*. Health monitoring was performed regularly under FELASA standards and samples taken analyzed by IDEXX Germany. Import and all experimental protocols were approved by the Icelandic Food and Veterinary Authority (license: 18081393)

### KMT2D variant testing

#### AlphaFold

For protein modeling and prediction, AlphaFold3 was used with the AlphaFold server, developed by Abramson et al. (Abramson et al., 2024). Input sequences were retrieved from UniProt (Bateman et al., 2017). KMT2D (UniProt ID: Q6PDK2), NCOA6 (UniProt ID: Q9JL19), PAXI1 (UniProt ID: Q6NZQ4), PGR1A (UniProt ID: Q99L02). Predictions were generated using the default parameters, with quality assessed by pLDDT scores. Structural visualization and analysis were conducted using ChimeraX-1.9 (Pettersen et al., 2021).

#### Gene expression

For *Kmt2d* expression, total RNA was extracted from 8-week-old hippocampus with TRIreagent (Thermo Fisher Scientific, AM9738), isolated with Direct-zol RNA MicroPrep kit (Zymo, R2060) and cDNA synthesis performed with High-Capacity cDNA Reverse Transcription Kit (Thermo Fisher Scientific, 4368814) according to manufacturer protocols. RT-qPCR was performed with Luna Universal qPCR Master Mix (NEB, M3003X) on a CFX96 Touch Real-Time PCR Detection System (Bio-Rad) using following primers: Kmt2d_E51-52_F (5’-AAACGGCCCCATACCTTGAACAGC-3’), Kmt2d_E52_R (5’-TGCACAAACTGCTTGCTGTACGGG-3’), Kmt2d_E49_F (5’-TCAGTGAGAACAATGGACGGCC-3’), (5’ATGATGCGGTTCCACACAGCTTGG-3’), Kmt2d_E2-3_F Kmt2d_E49-50_R (5’-AGATTCAGACCCAGCAGCTGATGG-3’), Kmt2d_E3_R (5’-ACACAGAATGGGAAGGTCTGGC-3’), Kmt2d_E3-4_F (5’-AAGCCTCCTCATGACTGCAGTAGG-3’), Kmt2d_E4_R (5’-AGATGGCAACTCAAAGCGCTGC-3’). Data processing was done with CFXmanager software (Bio-Rad).

#### Western Blot

Proteins were isolated from approximately 1mm of the spleen tissue, homogenized with traditional 1x Laemmli buffer followed by heating at 95°C for 5 minutes. KMT2D protein levels and H3K4 methylation quantification was performed by western blotting with 4–20% Mini-PROTEAN® TGX™ Precast Protein Gels (Bio-Rad, 4561093) and overnight transfer on PVDF membrane. Blocking was performed by 5%BSA in Tris Buffered Saline (TBS) with 0.01%Tween for about 4-6 hours at room temperature. Primary antibody staining was conducted overnight at 4°C with the following antibodies: 1:1000 KMT2D (MerckMillipore, ABE1867) with 1:1000 Vinculin (Sigma, V9131) as control or 1:1000 H3K4me1 (Abcam, ab8895), 1:1000 H3K4me3 (Abcam, ab8580) with 1:3000 H3 (Abcam, ab24834) as control. Secondary antibody staining was performed with IRDye® 800CW Donkey anti-Rabbit IgG (H + L) (LiCor, 926-32213) and IRDye® 680RD Donkey anti-Mouse IgG (H + L) (LiCor, 926-68072) and imaging using Odyssey CLx (LiCor).

### Mouse phenotyping

#### Growth

The body weight of each animal was measured weekly from 4- to 8-weeks postnatally. Analysis of skull anomalies were conducted on CT scans that were performed on 6-week old, euthanized mice. The CT system and pre-clinical optical imaging from MILabs (Houten, Netherlands) were used with the following setting: 50 kV tube voltage, 0.21 mA current with 1712 mGy dose estimate. Reconstruction was performed with MILabs software (v. 13.14) and images generated with Slicer (v. 5.6.2) (Fedorov et al., 2012). Long bones were isolated by dissection and measured at 3-4 months of age.

#### IgA blood serum levels

We assessed IgA levels in mouse blood serum samples using Enzyme-linked immunosorbent assay (ELISA). Submandibular blood collection was performed on 12-week-old mice to obtain serum. After collection, blood samples were allowed to coagulate at room temperature for 30 minutes, followed by centrifugation at 4000 rpm for 10 minutes to separate the serum from the pellet. The assay was conducted according to the manufacturer’s protocol for the Mouse IgA Uncoated ELISA Kit (Thermo Fisher, Catalogue number: 88-50450-88). Corning™ Costar™ 9018 ELISA plates were coated with 100 µL/well of capture antibody diluted in coating buffer. The plates were sealed and incubated overnight at 4°C. The following day, the wells were washed twice with 400 µL/well wash buffer. Thereafter, the wells were blocked with 250 µl of blocking buffer at room temperature for 2 hours. The wells were washed again as previously described. Standards and samples were prepared in 2-fold serial dilutions using assay buffer A (1X) and added to the wells. For blank wells, 100 µL/well of assay buffer (1X) was used. The plates were sealed and incubated for 2 hours at room temperature on a shaker at 400 rpm. The wells were washed, supplemented with 100 µL/well of detection antibody, sealed, and incubated on a shaker at 400 rpm at room temperature. After incubation, the washes were repeated for a total of 4 times. Subsequently, 100µL/well of substrate solution was added to each well and incubated for 15 minutes. The reaction was stopped by adding an equal volume of stop solution. The absorbance of the plates was measured at 450nm. One value was identified as significant outliers according to Grubbs’ test and was excluded from analysis.

#### Behavioral testing

All behavior testing was conducted sequentially, starting at about 8-10 weeks of age with Open field where mice were left to explore the open field (50×50cm) for 10 minutes and behavior tracked. Next, we performed Y-maze according to published protocol (Prieur & Jadavji, 2019). Briefly, mice were placed in a three-armed maze with one arm blocked for 5 minutes to explore followed by a trial with all arms available to the mice to explore. Measurements were made for the novel arm entries; the total time spent in the arm as well as calculations for the spontaneous alternations (%Spontaneous alternation=Alternations/Max alternations). The final behavior test conducted was Morris Water Maze with same setup as previously described (Bjornsson et al., 2014). A 150 cm diameter tank was filled with water that was allowed to equilibrate to room temperature and nontoxic white paint added. Mice were trained with 60 sec trials, first with visible platform testing at 4 trials per day for three consecutive days. Following a five-day training of 4-trials/day with a hidden platform. On the final day, the platform is removed, and the animal is allowed to swim for a total of 90 seconds. All tests were video recorded, measured and analyzed with Noldus equipment (Ethovision, version 14.0).

#### Immunofluorescence staining

After all behavior testing had been conducted, animals were injected with EdU over 7 consecutive days (50µg/g/day) followed by euthanasia with ketamine (16 mg/ml) and xylazine (4 mg/ml) mixture and perfusion with 4%PFA, 24-hours post last injection. Brains were fixed with 4%PFA for 24 hours followed by serial sucrose gradient (10% - 20% - 30%). Brains were then OCT embedded and cryosectioned. On the first day of immunostaining, tissue sections were washed twice with TBS-T (TBS; LiCor, 927–60001) containing 0.025% Triton X-100 (Sigma-Aldrich, T8787-100ML) for 10 minutes each. Antigen retrieval was performed by immersing the sections in sodium citrate buffer (pH 6) at 95°C for 20 minutes. Following antigen retrieval, sections were washed with 200 µL of 3% bovine serum albumin (BSA) prepared in PBS (pH 7.4) and subsequently incubated for 20 minutes with 200 µL of permeabilization solution (0.5% Triton X-100 in PBS). The sections were washed again with 3% BSA in PBS. Then, 100 µL of Click-iT™ Plus reaction cocktail, prepared according to the manufacturer’s protocol (Click-iT™ Plus EdU Imaging Kit; ThermoFisher Scientific, C10637), was added to each sample and incubated at room temperature for 1 hour. The sections were subsequently washed once with TBS-T. The sections were blocked with TBS-T containing 3% normal goat serum (NGS; Abcam, ab7481) for 1 hour at room temperature. Primary antibody, rabbit anti-Histone H3 (Tri-Methyl-K4) (Abcam, ab8580), was added to the blocking solution at a 1:300 dilution. The sections were incubated in this solution for 1 hour at room temperature, followed by overnight incubation at 4°C. The sections were washed three times with TBS-T, followed by incubation with secondary antibodies, goat anti-mouse IgG H&L (Alexa Fluor® 647; Abcam, ab150115) and goat anti-rabbit IgG H&L (Alexa Fluor® 555; Abcam, ab150078), diluted 1:1000 in blocking solution. The sections were incubated in this solution overnight at 4°C. After incubation, the sections were washed twice with TBS-T and stained with 1X Hoechst® 33342 solution (5 µg/mL; Click-iT™ Plus EdU Imaging Kit, ThermoFisher Scientific, C10637) for 1 hour at room temperature, protected from light. The sections were then washed two to three times with PBS, mounted using ProLong™ Gold Antifade Reagent (ThermoFisher Scientific, P36930), and stored at 4°C until imaging. Imaging was performed using a confocal microscope Cicero (CrestOptics) and an EVOS Auto microscope (ThermoFisher). Image preprocessing and quantification were conducted using ImageJ.

#### Histology staining

Intestinal Peyer’s patches were counted as previously described (Pilarowski et al., 2020). Kidneys were dissected from *Kmt2d^+/R5230H^*animals harboring either one or two kidneys as well as from *Kmt2d^+/+^*animals for histology comparison. H&E staining was performed according to protocol setup at the department of pathology, Landspitali – the National University Hospital of Iceland. Images of Hematoxylin &Eosin stained slides were captured with EVOS and glomeruli measured using ImageJ.

### Statistics

All plots and statistics were performed with GraphPad Prism (v.10.1.1) using either mixed-effects ANOVA or unpaired Student’s t tests. Significance levels were indicated with asterisks; *P ≤ 0.05, **P ≤ 0.01, ***P ≤ 0.001, ****P ≤ 0.0001.

---

**Supplementary Table 1.**
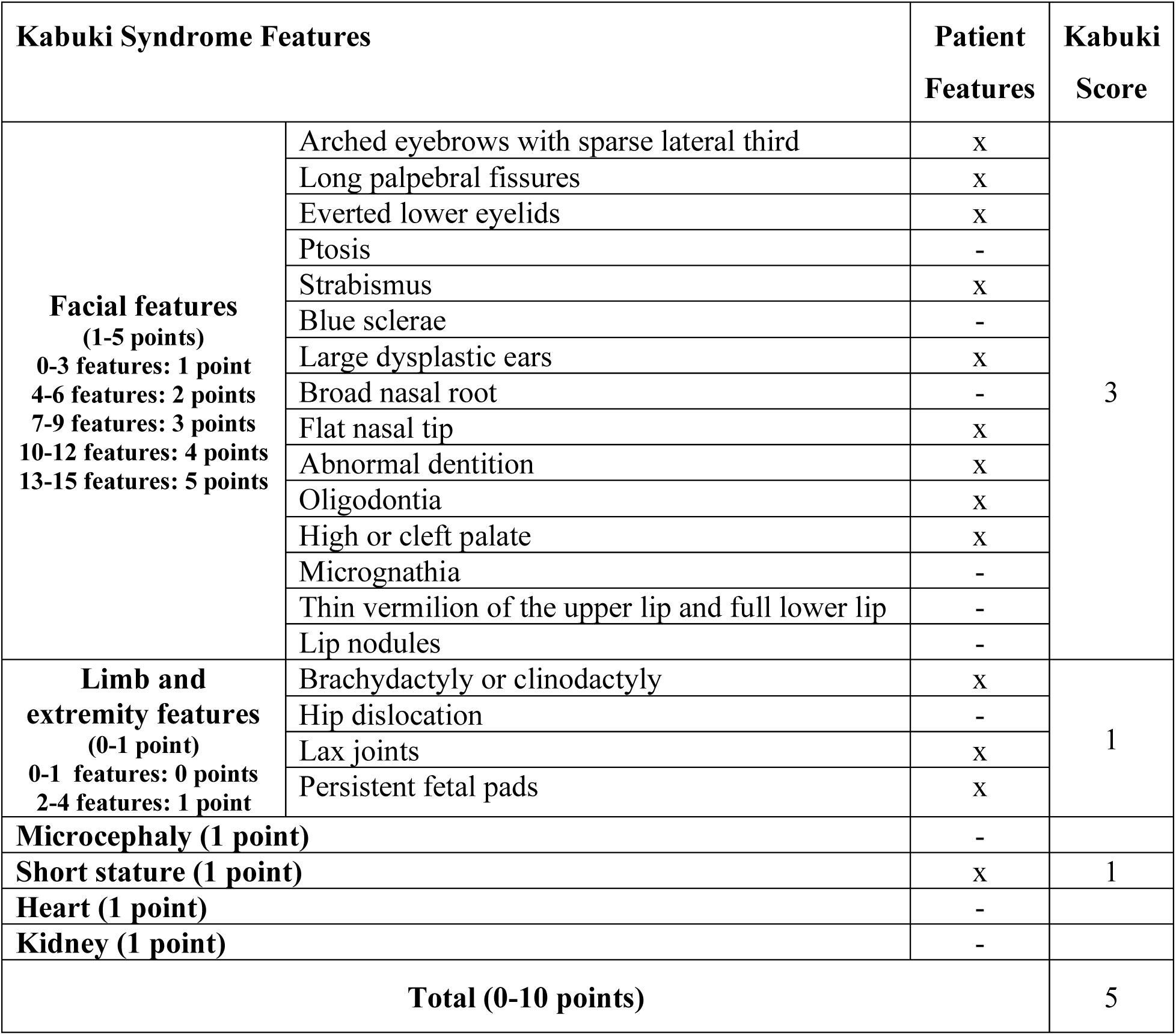
Symptoms presented in G15536A patient.

| **Kabuki Syndrome Features** |  | **Patient Features** | **Kabuki Score** |
| --- | --- | --- | --- |
| **Facial features**<br>**(1-5 points)**<br>**0-3 features: 1 point**<br>**4-6 features: 2 points**<br>**7-9 features: 3 points**<br>**10-12 features: 4 points**<br>**13-15 features: 5 points** | Arched eyebrows with sparse lateral third | x | 3 |
|  | Long palpebral fissures | x |  |
|  | Everted lower eyelids | x |  |
|  | Ptosis | - |  |
|  | Strabismus | x |  |
|  | Blue sclerae | - |  |
|  | Large dysplastic ears | x |  |
|  | Broad nasal root | - |  |
|  | Flat nasal tip | x |  |
|  | Abnormal dentition | x |  |
|  | Oligodontia | x |  |
|  | High or cleft palate | x |  |
|  | Micrognathia | - |  |
|  | Thin vermillion of the upper lip and full lower lip | - |  |
|  | Lip nodules | - |  |
| **Limb and extremity features**<br>**(0-1 point)**<br>**0-1 features: 0 points**<br>**2-4 features: 1 point** | Brachydactyly or clinodactyly | x | 1 |
|  | Hip dislocation | - |  |
|  | Lax joints | x |  |
|  | Persistent fetal pads | x |  |
| **Microcephaly (1 point)** |  | - |  |
| **Short stature (1 point)** |  | x | 1 |
| **Heart (1 point)** |  | - |  |
| **Kidney (1 point)** |  | - |  |
| **Total (0-10 points)** |  |  | **5** |

**Supplementary Figure 1.**
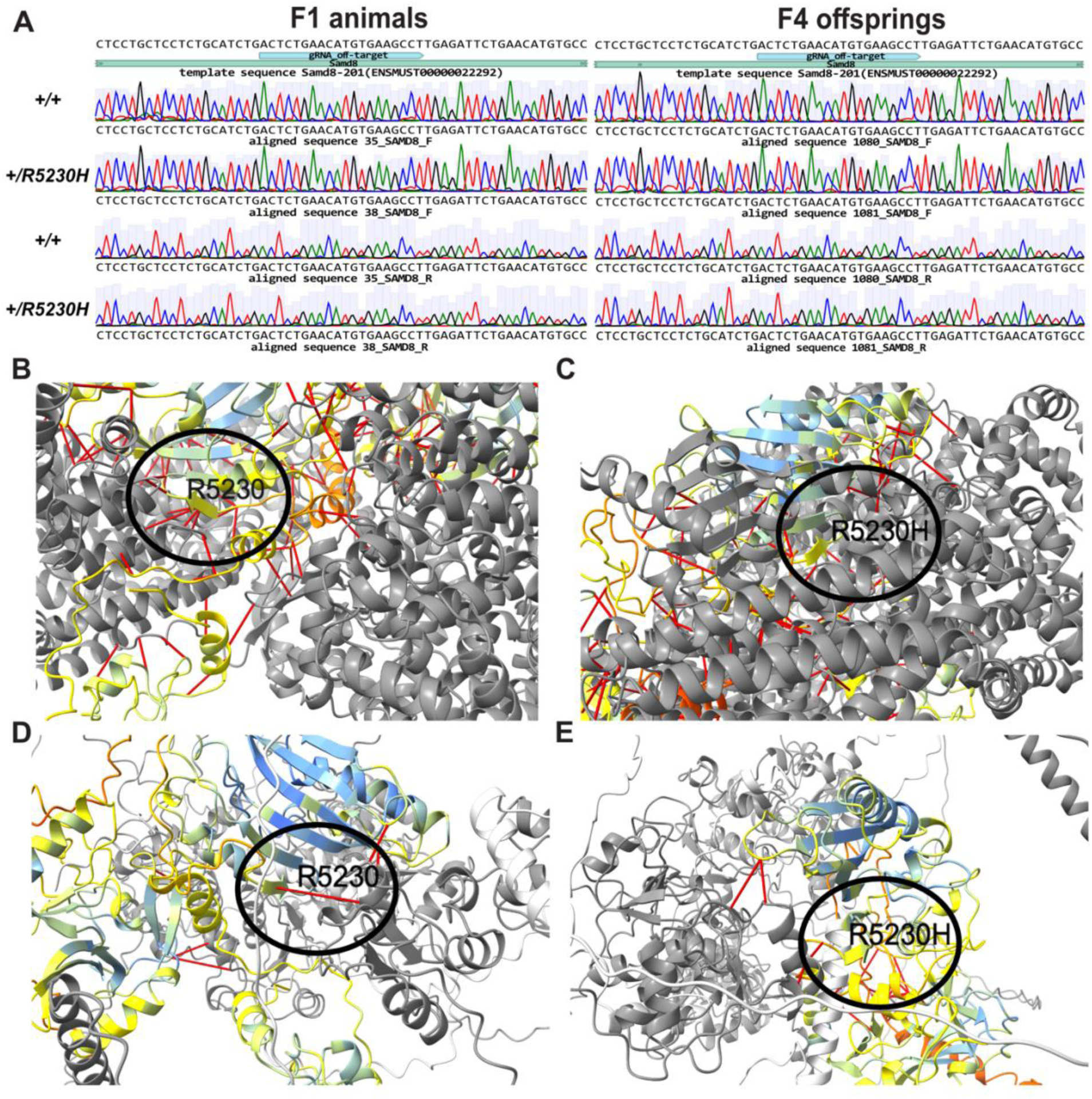
Predicted off-target and AlphaFold prediction of interaction with COMPASS components. Sanger sequencing results from CRISPR-Cas9 gRNA predicted off-target at *Samd8* loci in (A) F1 animals and F4 offsprings. AlphaFold3 prediction of part of KMT2D C-terminal starting at 100 amino acids downstream the FYRN domain. (B) Predicted interaction of NCOA6 colored in grey to R5230 or (C) R5230H KMT2D colored by pLDDT with circle showing interaction. (D) Predicted interaction of PAXI1 (grey) and its binding partner PGR1A (light grey) to R5230 or (E) R5230H KMT2D colored by pLDDT with circle showing interaction.

**Supplementary Figure 2.**
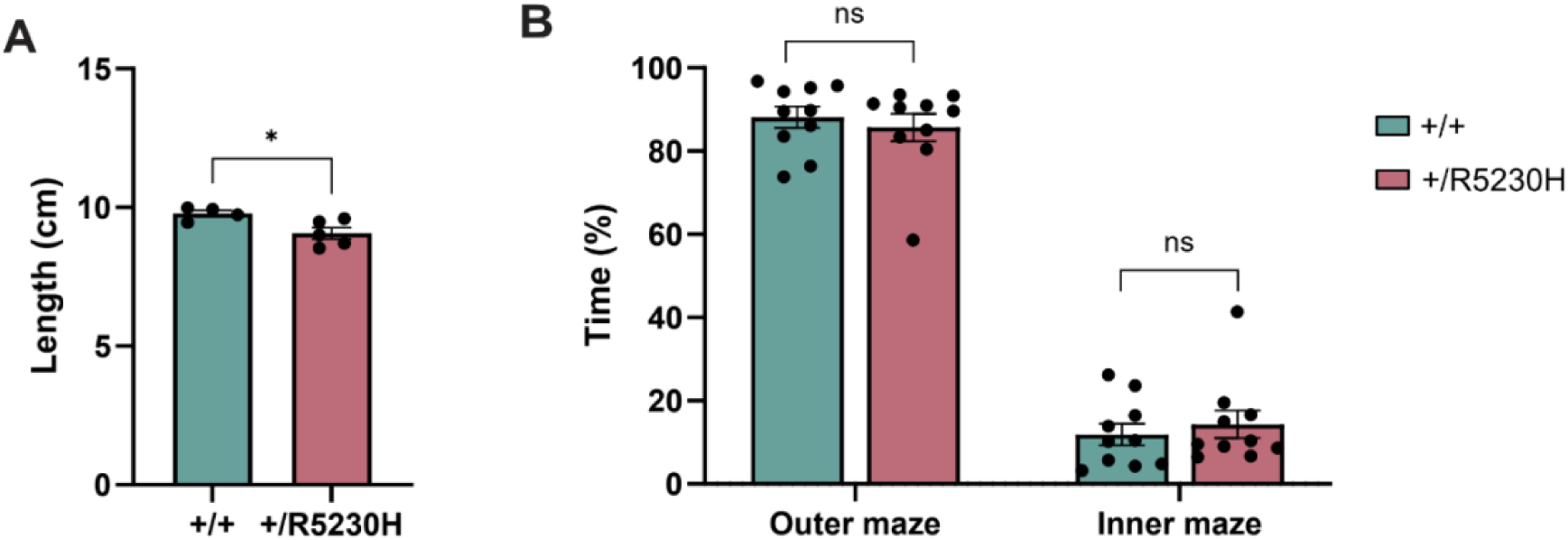
Phenotypes consistent across mouse models. (A) Body length measured in cm. (B) Open field test showing percentage of time spent in outer vs inner maze.

## Acknowledgements

We would like to thank Jon Gunnlaugur Jonasson and his team for performing pathological evaluation of kidneys and ArticLas staff members for performing the CT scans.

## Author contributions

H.T.B. and T.L. conceived the study. S.T.H., M.V., H.G. and E.D.B performed all experimental work. H.T.B., S.T.H., J.A.F., A.U. and M.V. interpreted the data. H.T.B. and S.T.H. wrote the initial draft of the paper. All authors edited the manuscript.

